# On the asymbiotic germination of some New Zealand spider orchids (*Corybas*)

**DOI:** 10.64898/2026.10.28.762650

**Authors:** Jennifer Alderton-Moss, Karin van der Walt, Carlos Lehnebach

## Abstract

*Corybas* is a large and widespread genus of terrestrial orchids, but little has been done to understand their germination. We tested germination of five species of helmet orchid (*C. cheesemanii, C. dienemus, C. iridescens, C. macranthus* and *C. oblongus*) on a range of asymbiotic media. *Corybas macranthus* germinated on four of seven tested media. From other species there was limited germination; for *C. cheesemanii* germination occurred for only two of five capsules collected concurrently and sown under the same conditions. The failure of filled seeds to germinate under optimal conditions could suggests the presence of seed dormancy. We trialed desiccation and warm moist stratification as methods to break dormancy, however this was unsuccessful.

## Introduction

The genus *Corybas* has ∼150 species (POWO, 2025), found through the Asia-Pacific region from the subantarctic islands to the Himalayas. In New Zealand (NZ), there are 22 species and a further 11 putative species (de Lange, et al., 2024). Of these, two species and two putative species are classified as Threatened, five species and four putative species are At Risk, and a further two species and two putative species are Data Deficient, meaning there is insufficient knowledge of population size and distribution to determine if they are threatened or not (de Lange, et al., 2024). Overseas, six of eight species studied on the Malay Peninsular are considered endangered (Go, et al., 2015); likewise, five of the six species found in China are endangered (Tao, et al., 2024). However, fewer than ten species have a current assessment for the IUCN red list (IUCN, 2025), reflecting wider gaps in threat assessments for plants (Gallagher, et al., 2023), particularly in places like New Guinea, which hosts approximately a third of all *Corybas* species (Lyon, 2016). Most *Corybas* are considered narrow endemics, which may make them more vulnerable to extinction, and specific threats include loss of habitat, particularly under predicted climate scenarios, and overcollection of plants for their ornamental value (Go, et al., 2015; Tao, et al., 2024). Developing conservation protocols, such as the ex-situ germination of *Corybas* seeds, should thus be prioritised.

Like all orchids, *Corybas* have dust seeds that do not contain the required nutriment to support their germination (Brundrett & Tedersoo, 2018). Orchid germination is facilitated by a relationship between the orchid and a mycorrhizal fungus, often in the fungal groups Ceratobasidiaceae, Sebacinales and Tulasnellaceae (Dearnaley, et al., 2016). Indeed, all these groups have been found to associate with mature plants of the genus *Corybas* (Alderton-Moss, et al., 2024; Lyon, 2014; Watkins, 2012; Qin, et al., 2024).

Conservation of threatened orchids can be achieved by inoculating seed with a mycorrhizal fungus and germinating seed *ex situ*, a process known as symbiotic germination. Attempts of the symbiotic germination of *Corybas iridescens* with various fungi is reported by Watkins (2012), however protocorm formation was not achieved. Additionally, we conducted a pilot study into the symbiotic germination of eight *Corybas* spp. in NZ; although germination occurred, it was noted that protocorm development for many species stalled after this initial germination (Alderton-Moss, et al., 2024). For certain species, it has also been noted that the fungus grows very slowly (Warcup, 1981) or culturing has failed entirely (Lyon, 2014; Alderton-Moss, et al., 2023), adding further challenges to the symbiotic germination approach.

Alternatively, orchids may be propagated using an asymbiotic approach, in which media contain all the nutrients required for the germination of the orchid. This approach, which eliminates the requirement for a fungus, has proved successful for germinating many terrestrial orchids, including some NZ species (van der Walt, et al., 2024; Diantina, et al., 2020). However, there is little published research into the asymbiotic germination of *Corybas*. Germination of several species has been reported to occur on Knudson C medium (Alderton-Moss, et al., 2024), however protocorm development occurred very slowly, suggesting suboptimal germination conditions. A variety of commercial asymbiotic media are available, and different orchid species seem to have specific media preferences (Dowling & Jusaitis, 2012; Van Waes & Debergh, 1986; Kauth, et al., 2008).

Germination efforts may be further complicated by seed dormancy, defined as “the failure of seeds to germinate although environmental conditions including water, temperature, light and gases are favorable for germination” (Baskin & Baskin, 2014). Dormancy in orchid seeds has long been recognised (Knudson, 1950) and may be due in part due to an impermeability of the seed coat (Kauth, et al., 2008; Rasmussen, 1995). This is particularly noticeable in terrestrial species (Rasmussen, 1995). Preliminary germination studies into *Corybas* reported germination from only one of two *C. macranthus* capsules when seeds were sown under the same conditions, suggesting the presence of seed dormancy in *Corybas* (Alderton-Moss, et al., 2024). Effective germination of *Corybas* may thus require dormancy breaking. Several strategies have previously been employed for orchids, including desiccation (Batty, et al., 2001), cold stratification (Rasmussen, 1995; Baskin & Baskin, 2014), and warm stratification (Janissen, et al., 2022; Rasmussen, 1995). As yet, there have been no reports of dormancy breaking treatments used for *Corybas*.

Conservation outcomes of *Corybas*, both in NZ and abroad, can be improved by understanding how to effectively propagate them. In this study we aim to begin addressing the knowledge gaps surrounding *Corybas* germination. Specifically, we intend to 1) compare various asymbiotic media to determine which media are most favorable for *Corybas* germination; 2) compare germination across capsules and determine if seed germinability correlates to capsule features; and 3) determine if dormancy breaking methods improve or facilitate germination.

## Methods

### Capsule collection

*Corybas* capsules were collected prior to dehiscence from the Wellington region (lower North Island, NZ), between August 2024 and January 2025. These included *C. cheesemanii* (n=8, Figure 1A), *C. dienemus* (n=1, Figure 1B), *C. iridescens* (n=1, Figure 1D), *C. macranthus* (n=4, Figure 1E) and *C. oblongus* (n=2, Figure 1F). Capsules were surface sterilised in 70% propanol for 2 minutes, air dried in a laminar flow cabinet, and cut open to sow seed for germination trials. Germination was set up within 24 hours of collection, unless otherwise stated.

**Figure 1.**
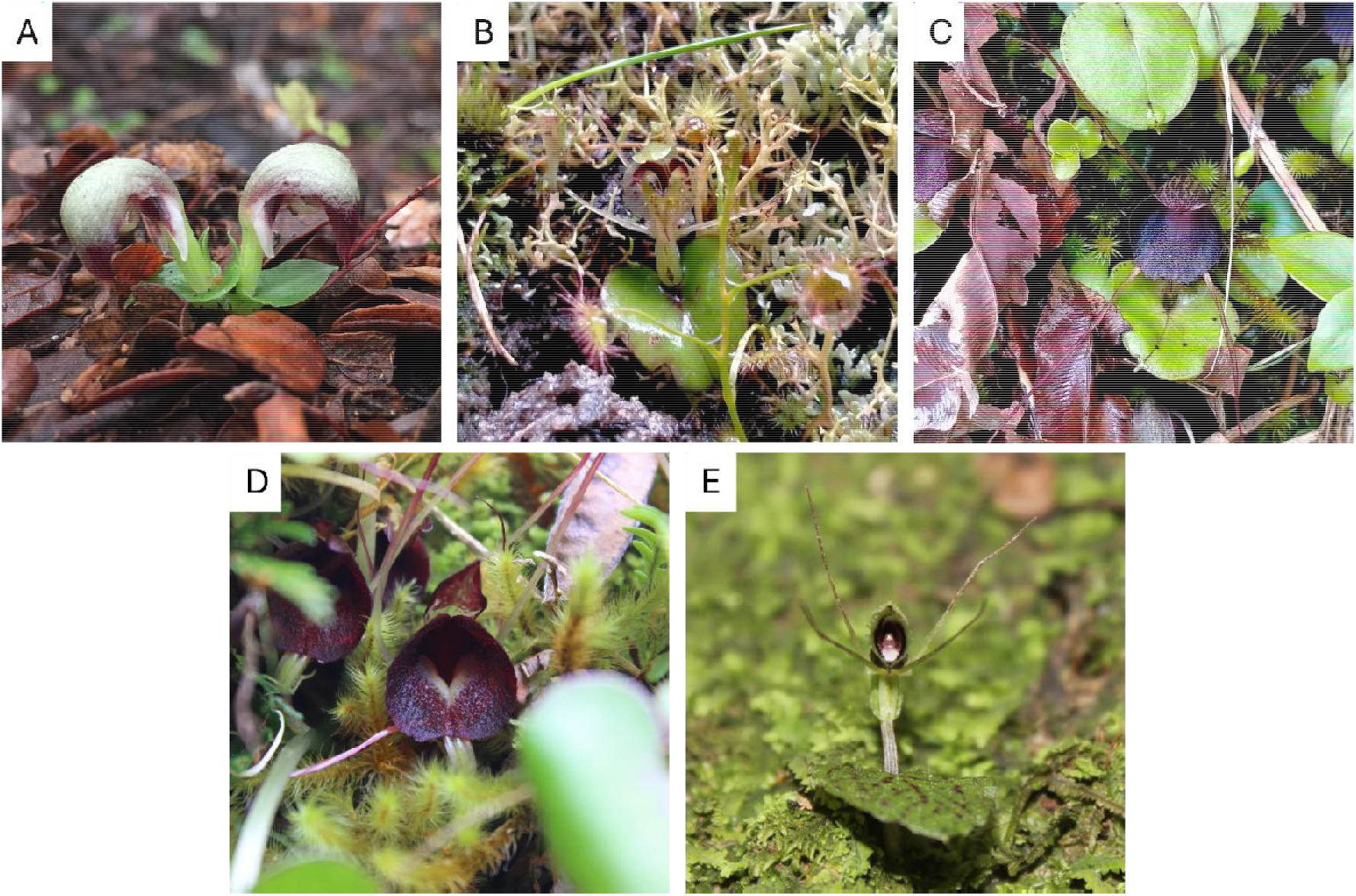
New Zealand *Corybas* species included in the present study: A) *C. cheesemanii*, B) *C. dienemus*, C) *C. iridescens*, D) *C. macranthus,* and E) *C. oblongus*.

### Asymbiotic media test

To identify the optimal media for asymbiotic germination of *Corybas* spp., seed was sown on a range of commercially available, asymbiotic media. The following media were included in the test: BM1, Knudson C Morel Modification (KC), Modified Melins Norkrans with sucrose (MMN+suc), Murashige and Skoog with sucrose (MS+suc), P723, T839 and Vacin and Went with sucrose (VW+suc). Agar and sucrose levels were normalised across media to 10g/L and 20g/L, respectively. Seed of five *Corybas* species (i.e. *C. cheesemanii*, *C. dienemus*, *C. iridescens*, *C. macranthus* and *C. oblongus*) were sown on each media and incubated in the dark at 12/25 °C. Germination was monitored every 4 weeks and scored after 24 weeks. A single capsule from each species was used for this trial. Although between 5 and 8 reps were initially plated for each species and each media, due to contamination *C. dienemus* and *C. cheesemanii* on VW+suc were discarded. Additionally, *C. iridescens* was not sown on MS+suc, and *C. cheesemanii* was not sown on P723 or T839.

### Capsule germinability test

Being able to identify capsules *in situ* where seed will readily germinate will aid the propagation of *Corybas* species. The following capsule features of *C. macranthus* and *C. cheesemanii* were recorded: capsule width, capsule length, peduncle length, and capsule maturity. Capsule measurements were made under stereomicroscope using cellSens software; peduncle length was measured using a caliper. Capsule maturity was based off capsule colour (light green for immature, green-brown for mature) and seed coat colour (white for immature, brown for mature). As germination on KC had previously been observed for *C. cheesemanii* and *C. macranthus* (Alderton-Moss, et al., 2024), seed from all capsules was sown on KC. For two immature capsules of *Corybas cheesemanii*, seed was also sown on BM1. Germination, defined as the development of rhiozids, was monitored every 4 weeks and scored after 24 weeks.

### Dormancy breaking test

Based on the observation that germination of *Corybas* spp. varies between capsules, we trialed some common dormancy breaking methods. Fresh seed from mature capsules of *C. cheesemanii* and *C. macranthus* underwent the following treatments: desiccation (24 hours over silica gel followed by sowing on KC), warm moist stratification (seed sown on KC, incubated at 25 °C for one week), and desiccation followed by warm moist stratification (dried 24 hours silica gel, sown on KC, incubated at 25 °C for one week). Warm moist stratification was chosen as *C. cheesemanii* releases seed in spring and *C. macranthus* in early summer, suggesting seeds must survive the summer before germination (see studies on *Caladenia* reported in Janissen, et al., 2022). Some seeds were left untreated as a control. Seed was sown on KC, and germination was scored after 24 weeks.

## Results

### Asymbiotic media test

*Corybas macranthus* germinated on four of the seven tested media: BM1 (11.4%), MMN+Suc (2.3%), MS+Suc (3.5%) and VW+Suc (7.1%) after 24 weeks (Figure 2). Seed germinationwas observed on BM1 after 8 weeks, but not until 16 weeks on VW+Suc and MMN+Suc, and 24 weeks on MS+Suc. Although a greater proportion of seed germinated on BM1 compared to the other media, and germination occurred more quickly, no significant difference was found at 24 weeks between germination on BM1, MMN+Suc, MS+Suc and VW+Suc. Only a single *Corybas iridescens* seed germinated on P723, and for *C. cheesemanii*, *C. dienemus* and *C. oblongus* no germination was observed in the present trial (Table 1).

**Figure 2.**
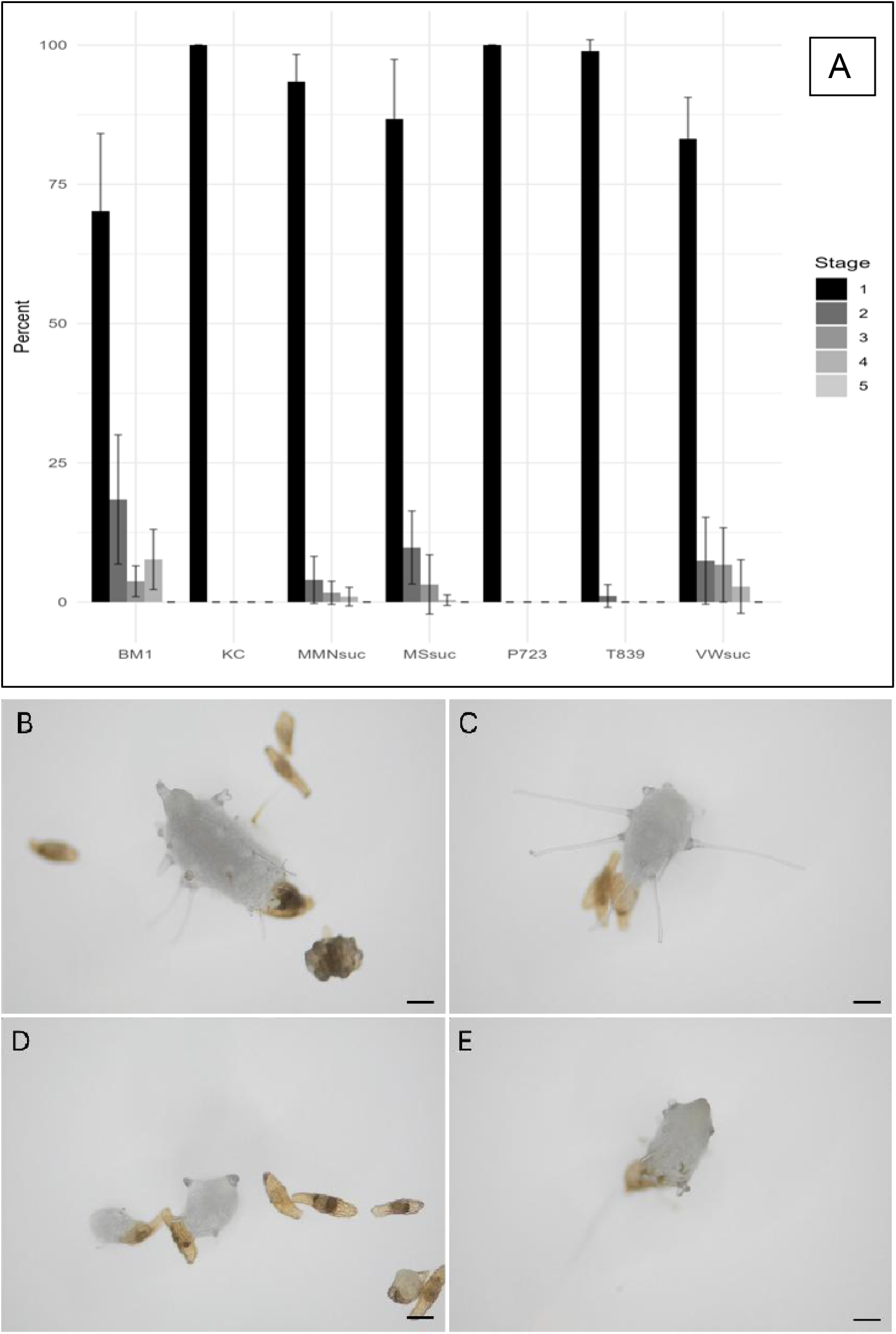
A) Percent germination of *C. macranthus* on asymbiotic media and developing protocorms after 24 weeks on B) BM1, C) MMN+Suc, D) MS+Suc, and E) VW+Suc. Scale bars are 200μm.

**Table 1.** Germination of each *Corybas* species after 24 weeks on the tested asymbiotic media. Seed/media combinations that resulted in germination are indicated by ‘+’ and that did not result in germination are indicated by ‘–’. The * indicates that this seed/media combination resulted in germination in a previous trial (Alderton-Moss et al. 2004).

|  | BM1 | KC | MMN+suc | MS+suc | P723 | T839 | VW+suc |
| --- | --- | --- | --- | --- | --- | --- | --- |
| *Corybas cheesemanii* | —* | —* | — | — |  |  |  |
| *Corybas dienemus* | — | — | — | — | — | — |  |
| *Corybas iridescens* | — | —* | — |  | + | — | — |
| *Corybas macranthus* | + | —* | + | + | — | — | + |
| *Corybas oblongus* | — | —* | — | —* | —* | — | — |

### Capsule germinability test

Germination rates of *C. cheesemanii* capsules collected at the same time and treated under the same conditions showed vastly different abilities to germinate (Table 2). For *C. cheesemanii*, of the eight capsules collected, only two germinated on KC, Cap24d (0.8% germination, Figure 3c) and Cap24e (6.8% germination, Figure 3d). Additionally, two immature capsules with underdeveloped white seed germinated on BM1 but not KC (Figure 3a and 3b). As these did not germinate on KC, it supports the idea that other media may be more suitable, at least for immature seeds. Capsule germinability for *C. macranthus* could not be determined as no seed germinated on KC (Table 2). Although Cap24a germinated on other media (BM1, MMN+Suc, MS+Suc, VW+Suc), seed from the other capsules was only tested on KC, thus the lack of germination may be due to unsuitable growing conditions rather than poor germinability of the seed.

**Figure 3.**
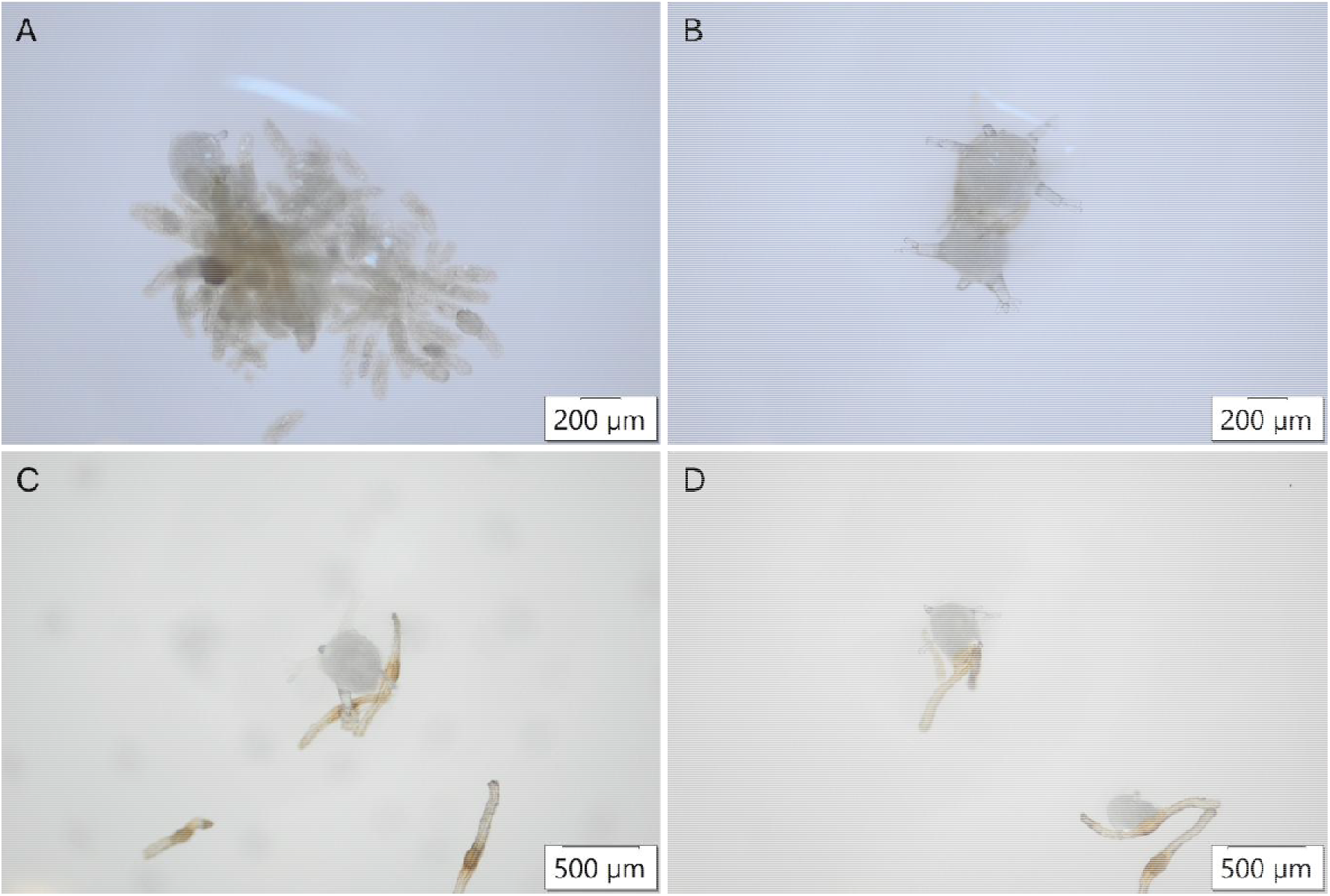
*Corybas cheesemanii* seed germinating on BM1 from Cap24a (A) and cap 24b (B) and on KC from Cap24d (C) and Cap24e (D).

**Table 2.**
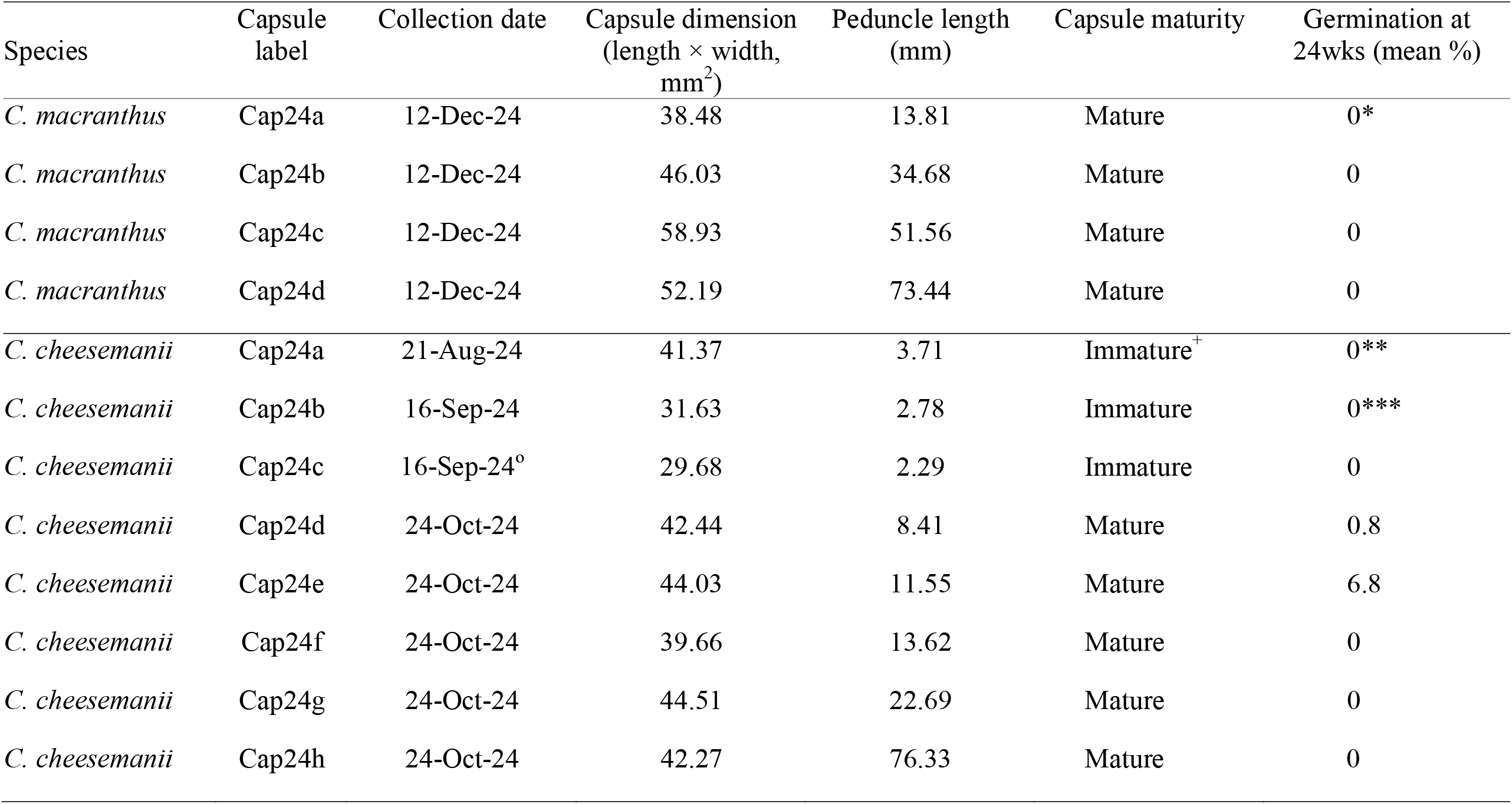
Morphological features recorded for each capsule and germination data after 24 weeks on KC and BM1 (Cap24a and CAP24b, only). *Seed germinated on BM1 (11.4%), MMN+Suc (2.3%), MS+Suc (3.5%) and VW+Suc (7.1%). **Seed germinated on BM1 (4.4%; seed with no embryo were not included in this calculation). ***Seed germinated on BM1 (79.5%; seed with no embryo were not included in this calculation). ^+^Most seed lacking embryo. °Capsule not sown until 30-Sep-24 and had begun to decay before seed sowing.

### Dormancy breaking test

For *Corybas cheesemanii* and *C. macranthus,* seeds from capsules used for the dormancy breaking trial failed to germinate regardless of whether they were subjected to desiccation, warm moist stratification, desiccation followed by warm moist stratification, or no treatment.

## Discussion

### Asymbiotic media test

The media preference displayed by *C. macranthus* informs future asymbiotic germination of *Corybas*, however our understanding is limited by the lack of germination from the other *Corybas* species. Interestingly, in this trial no seed germinated on KC, despite previous findings that *C. macranthus* can germinate on KC (Alderton-Moss, et al., 2024). Previous studies have shown that different orchid species will have different preferences for asymbiotic media (Kauth, et al., 2008; Van Waes & Debergh, 1986; Dowling & Jusaitis, 2012). This observation can extend to congeneric orchids, for example, Diantina *et al*. (2020) reported that different *Dendrobium* species showed different preferences to asymbiotic media. If asymbiotic media is mimicking the role of a mycorrhizal fungus, it may be expected that orchids that associate with different fungal taxa, or who occupy different habitats, would thus have different preferences in asymbiotic media. This may be particularly relevant in *Corybas*, where different species of fungi are reported for different species/clades/regions (e.g., Serendipitaceae is reported in China (Qin, et al., 2024) and Tulasnellaceae in Australia (Lyon, 2014)), and where habitats in NZ range from forest understory to open wetlands. Further work is needed to determine if other *Corybas* share the preferences of *C. macranthus*, or if there are species specific media preferences.

### Capsule germinability test

Rasmussen (1995) reports that germination happens more readily for immature seeds of terrestrial orchids, including for some species where germination of mature seeds could not be achieved, likely due to a dormancy that is acquired as seeds develop. For example, for several species of the terrestrial orchid genus *Cypripedium*, germination rates were best with seeds that were slightly immature, having an early globular embryo smaller than that of mature seeds (Lee, et al., 2005; Zhang, et al., 2013; De Pauw & Remphrey, 1993). Interestingly, both seed that was collected earlier in the ripening process and seed that was collected later was less likely to germinate, although the optimal time for seed collection differed between species. There may be several barriers to germination of mature orchid seeds, including hydrophobicity of the testa (Rasmussen, 1995) or higher levels of germination inhibitors such as abscisic acid (Van der Kinderen, 1987). There have previously been no studies exploring the germinability of mature versus immature seeds of *Corybas*.

Generally, *Corybas* flowers develop on a short peduncle which sits just above their solitary leaf; this short stem elongates as the fruit matures, presumably so that the capsule is better situated for wind dispersal of seeds (Moore & Edgar, 1970). Watkins (2012) monitored peduncle extension for three specimens of *Corybas iridescens* and found that peduncles were fully extended 63±9 days after pollination, with extension accelerating in the final ten days prior to dehiscence. Peduncle length may thus serve as a proxy for capsule maturity, allowing for the selection of immature capsules with ready to germinate seed, without requiring timed hand-pollination events and repeated site visits. Our finding in the present study with *C. cheesemanii* supports the idea that capsules with shorter peduncles may germinate more readily, including where the seed has not fully matured. Additionally, while germination did not occur from mature *C. oblongus* capsules collected late January 2025 (Table 1), seed from immature capsules collected in late December 2024 did germinate. Unfortunately, due to the clumped nature of the immature seed, germination could not be quantitatively scored. Although our study supports the idea, the limited sample sizes make it far from conclusive; we recommend further research that directly links time after pollination, peduncle length and seed germinability.

### Dormancy breaking test

The dormancy breaking methods trialed in the present study did not promote germination of *Corybas,* however this may have been due to suboptimal germination medium and we recommend repeating experiments on BM1. We hypothesized that *Corybas* seeds must be desiccated prior to germination, as they are typically released late spring/summer when they would be subject to seasonal drying. Likewise, we hypothesized that *Corybas* must be subjected to warm stratification due to the seeds being released from the end of spring until mid-summer (see studies on *Caladenia* reported in Janissen, et al., 2022), however it is currently unknown when *Corybas* germinate *in situ*. Identifying the time of germination would provide insights into possible environmental triggers for dormancy and germination, however it may prove very challenging to locate protocorms.

## Conclusions

While we are excited to report on the asymbiotic germination of *Corybas cheesemanii* and *C. macranthus*, there is still considerable work to do to overcome the low germination rates, inconsistent results, and failure for other species of *Corybas* to germinate under comparable conditions. We also acknowledge that although we have observed germination and protocorm formation, these protocorms are yet to be transitioned out of sterile culture and are unlikely to be of a size that would survive *in situ* or *ex situ* in a nursery. We recommend further investigation into the possible dormancy of *Corybas* seeds, as well as how to optimise germination conditions to better support their germination and continued growth.

## Notes

### Competing Interest Statement

The authors have declared no competing interest.

### Funding

- Australian Orchid Foundation: Project 356/2023

